# Deletion of *Fibroblast growth factor 9* globally and in skeletal muscle results in enlarged tuberosities at sites of deltoid tendon attachments

**DOI:** 10.1101/2021.02.23.432498

**Authors:** Connor C. Leek, Jaclyn M. Soulas, Iman Bhattacharya, Elahe Ganji, Ryan C. Locke, Megan C. Smith, Jaysheel D. Bhavsar, Shawn W. Polson, David M. Ornitz, Megan L. Killian

## Abstract

The growth of most bony tuberosities, like the deltoid tuberosity (DT), rely on the transmission of muscle forces at the tendon-bone attachment during skeletal growth. Tuberosities distribute muscle forces and provide mechanical leverage at attachment sites for joint stability and mobility. The genetic factors that regulate tuberosity growth remain largely unknown. In mouse embryos with global deletion of *fibroblast growth factor 9* (*Fgf9*), the DT size is notably enlarged. In this study, we explored the tissue-specific regulation of DT size using both global and targeted deletion of *Fgf9*. We showed that cell hypertrophy and mineralization dynamics of the DT, as well as transcriptional signatures from skeletal muscle but not bone, were influenced by the global loss of *Fgf9*. Loss of *Fgf9* during embryonic growth led to increased chondrocyte hypertrophy and reduced cell proliferation at the DT attachment site. This endured hypertrophy and limited proliferation may explain the abnormal mineralization patterns and locally dysregulated expression of markers of endochondral development in *Fgf9^null^* attachments. We then showed that targeted deletion of *Fgf9* in skeletal muscle leads to postnatal enlargement of the DT. Taken together, we discovered that *Fgf9* may play an influential role in muscle-bone crosstalk during embryonic and postnatal development.

## Introduction

The transmission of muscle forces to bone via tendons is required for the functional growth of the vertebrate skeleton (Arvind and Huang, 2017; Biewener et al., 1996; Blitz et al., 2009; Kahn et al., 2009; Rot-Nikcevic et al., 2006). Bone ridges and tuberosities are skeletal “superstructures” that transmit muscle forces to the periosteal surface of long bones and increase leverage at sites of tendon attachments (Benjamin et al., 2006; Blitz et al., 2009; Blitz et al., 2013; Zelzer et al., 2014). The presence of superstructures requires both initiation of a modular *Sox9*+/*Scx*+ progenitor cell pool (Blitz et al., 2013) and maintenance of skeletal muscle contraction (Blitz et al., 2009; Hamburger and Waugh, 1940; Pai, 1965; Rot-Nikcevic et al., 2006; Tremblay et al., 1998). One of the most prominent superstructures in the mouse skeleton is the deltoid tuberosity (DT) of the humerus, which is a migratory tendon attachment that projects anterolaterally from the shoulder with a fin-like shape (Felsenthal et al., 2018). Migratory tendon attachments are first established by a population of *Sox9*+ progenitor cells during embryonic development. Subsequently, these *Sox9*+ cells are then replaced by *Gli1*+ progenitor cells during postnatal growth (Felsenthal et al., 2018). As such, the growth of tendon attachments is likened to that of an arrested growth plate (Benjamin et al., 2006; Schwartz et al., 2015), with the attachment includes a layered arrangement of hypertrophic and pre-hypertrophic *Gli1+* cells, similar to the growth plate during endochondral ossification, as well as a mineralized gradient at the enthesis between tendon and bone (Benjamin et al., 2006; Schwartz et al., 2015; Zelzer et al., 2014).

Because it is a site of tendon attachments, the DT is also a phenotypic readout for estimating muscle loads during embryonic development in mice. For example, in muscular-compromised mouse models (*Myf5^null^*: *MyoD^null^*, *Mdg*, and *Spd* mutants), the DT fails to maintain its size and shape during the mid-to-late stages of embryonic growth (Blitz et al., 2009; Hamburger and Waugh, 1940; Pai, 1965; Rot-Nikcevic et al., 2006; Tremblay et al., 1998). Conversely, mice with genetic mutations that lead to increased muscle mass (e.g., *myostatin^-^*^/-^ mice) exhibit an autonomous increase in the size and shape of the DT (Hamrick et al., 2002). Though many mutations related to skeletal muscle unloading result in smaller bone ridges (Blitz et al., 2009; Hamburger and Waugh, 1940; Pai, 1965; Rot-Nikcevic et al., 2006; Tremblay et al., 1998), embryos that lack the protein-encoding gene for Fibroblast growth factor 9 (*Fgf9*) have enlarged DTs (Hung et al., 2007). Surprisingly, even though the DTs are enlarged, the stylopods (i.e., humeri and femurs) of *Fgf9* mutant (*Fgf9^null^*) embryos are shorter and experience delayed growth plate hypertrophy and proliferation, as well as reduced angiogenesis, ultimately restricting osteogenesis during long bone development (Hung et al., 2007; Hung et al., 2016). These phenotypic differences between DT growth and long bone development in *Fgf9^null^* embryos suggest a unique role of *Fgf9* as a negative regulator of growth in attachment superstructures.

FGF9 is one of three ligands in the fibroblast growth factor (FGF) family known to be involved in bone development (Berendsen and Olsen, 2015; Coutu et al., 2011; Deng et al., 1996; Jacob et al., 2006; Karuppaiah et al., 2016; Ohbayashi et al., 2002; Ornitz and Itoh, 2015; Ornitz and Marie, 2002; Ornitz and Marie, 2015; Sahni et al., 1999; Su et al., 2014; Vaahtokari et al., 1996). In the developing limb, *Fgf9* is expressed as early as embryonic day 10.5 (E10.5) within the apical ectodermal ridge (Colvin et al., 1999; Garofalo et al., 1999; Hung et al., 2007; Yang and Kozin, 2009). As limb development progresses, expression of *Fgf9* coincides with expression of other chondrogenic markers, such as *Fgfr2* and *Fgf18* (Govindarajan and Overbeek, 2006; Hung et al., 2007; Lu et al., 2015). In periosteal and trabecular osteoblasts, overexpression of *Fgf9* leads to smaller mice with shorter growth plates (Karuppaiah et al., 2016). Although *Fgf9* expression is required for long bone growth, its expression is predominantly found in the surrounding muscle and perichondral cells and not in the bone anlagen, suggesting that FGF9 may also originate from surrounding soft tissues (Colvin et al., 1999; Garofalo et al., 1999; Hung et al., 2007). However, the role of tissue specific *Fgf9* on the growth of bony superstructures like the DT has not been explored.

In this study, we hypothesized that global knockout of *Fgf9* leads to increased expansion and growth of the arrested growth plate at sites of migratory tendon attachments. We quantified the size of the DT superstructure using whole mount staining and characterized cell shape and size using histomorphometry. We then used multiplexed fluorescence *in situ* hybridization (FISH) at key timepoints during embryonic growth to visualized expression of growth plate markers at the tendon attachment of the DT in wildtype and *Fgf9^null^* limbs. We used RNA sequencing (RNAseq) to identify differentially expressed genes between WT and *Fgf9^null^* bone anlagen and skeletal muscle during late embryonic development. Surprisingly, we found significant differential expression in muscle, but not bone. Therefore, we developed conditional muscle-specific *Fgf9* knockout mice (*Fgf9^cKO^*) to determine the effect of muscle-specific deletion of *Fgf9* on postnatal DT size and skeletal muscle innervation. Our findings suggest a muscle-specific role of FGF9 in the size of stylopod superstructures, further expanding the role of FGF signaling in musculoskeletal development.

## Results

### Global loss of *Fgf9* results in an enlarged DT, even with a shortened humerus

To confirm the larger size of the DT in *Fgf9^null^* neonates suggested previously by Hung et al., we used whole-mount staining to visualize cartilage with Alcian Blue and mineralized bone with Alizarin Red (Hung et al., 2007). We found that *Fgf9^null^* neonates had a 68% larger ratio of the DT area to total humerus area compared to WT neonates at postnatal day 0 (P0) (Figure 1A,B). *Fgf9^null^* neonates also had a similar total humerus area compared to WT neonates (Supplemental Figure 2B). We did not find a sex-dependent effect of global *Fgf9* deletion on DT size (Figure 1, blue and pink dots).

**Figure 1:**
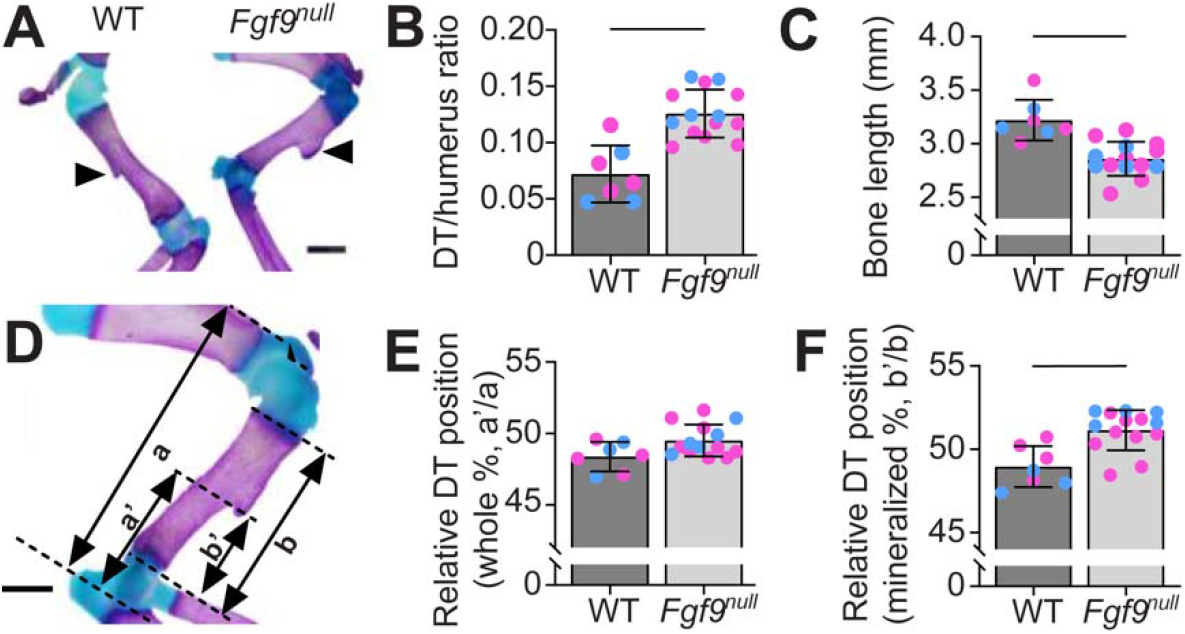
The deltoid tuberosities (DTs) of *Fgf9* global knockout (*Fgf9^null^*) neonates were larger compared to WT neonates. **(A)** Whole mount-stained forearms of a (left) WT mouse and (right) *Fgf9^null^* neonate at postnatal day 0 (P0) (Scale bar = 1mm). Black arrows point to the DT, which was noticeably larger on *Fgf9^null^* neonate humeri. Alcian Blue stain was present on the DT of *Fgf9^null^* neonates but absent in WT neonates. **(B)** Normalized area of the DT was significantly larger in *Fgf9^null^* neonates compared to WT neonates at P0. Whether the mice were male (blue) or female (pink) had no effect. **(C)** Contrastingly, the enlarged DT was a part of a shorter mineralized humerus. **(D)** The positioning of the DT on the humerus with respect to the total length (a, a’) and the mineralized length (b, b’). **(E)** Relative to the whole humeri, the DT of *Fgf9^null^* neonates was in a similar relative location compared to WT neonates. **(F)** Relative to the mineralized region, the DT of *Fgf9^null^* neonates was more proximally placed than WT neonates. (Data are biological replicates, mean ± SD. Lines in B, C, and F indicate significance (p < 0.05).

We observed increased Alcian Blue-stained cartilage on the crest of the DT in *Fgf9^null^* neonates, but this cartilage was absent on the crest of embryonic stage-matched WT DTs (Figure 1A). In line with previously published work, *Fgf9^null^* neonates had shorter humeri (Figure 1C) than WT neonates at P0 (*Fgf9^null^* = 3.2 ± 0.2mm, WT = 2.9 ± 0.2 mm) (Hung et al., 2007). DT migration was measured relative to the length of the humerus (Figure 1D). In *Fgf9^null^* neonates, the DT was located in a relatively similar position on the humerus compared to WT littermates (Figure 1E,F). Using dynamic bone histomorphometry, *Fgf9^null^* DTs had an increased and more variable mineral growth rate compared to WT embryos (mineral apposition rate between E16.5 and E18.5: *Fgf9^null^* = 133.1 ± 117.7%, WT = 32.9 ± 10.7%).

### Global loss of *Fgf9* led to larger attachment-site chondrocyte size but reduced cell proliferation at the attachment

Next, we compared the cellular and matrix morphology of the DT attachment between WT and *Fgf9^null^* embryos using Hematoxylin and Eosin Y (H&E) staining of transverse histological sections. At E15.5, the size and cell morphology of the DT was similar between WT and *Fgf9^null^* embryos (Figure 2A,E). At E16.5, the DT of *Fgf9^null^* embryos appeared larger and less mineralized compared to the DT of WT embryos (Figure 2B,F). At E18.5 and P0, we observed more hypertrophic chondrocytes in the DT of *Fgf9^null^* mice (Figure 2G,H) compared to WT mice (Figure 2C,D). These hypertrophic chondrocytes stained positive for type X collagen at E16.5 samples (Supplemental Figure 3). At E16.5, we measured a similar number of hypertrophic chondrocytes at the DT for *Fgf9^null^* and WT littermates (Figure 2I). However, the average size of DT-localized hypertrophic chondrocytes in *Fgf9^null^* embryos at E16.5 was 72% larger compared to WT embryos (Figure 2L). Additionally, cell proliferation at E16.5, visualized using 5-ethynyl-2’-deoxyuridine (EdU), was 51% lower in attachments but not tendons or perichondrium, of *Fgf9^null^* embryos compared to similar regions of WT embryos (Figure 2N). We did not find measurable differences in cell density (Hoechst-stained; nuclei/100μm^2^) between E16.5 WT and *Fgf9^null^* embryos (Figure 2K).

**Figure 2:**
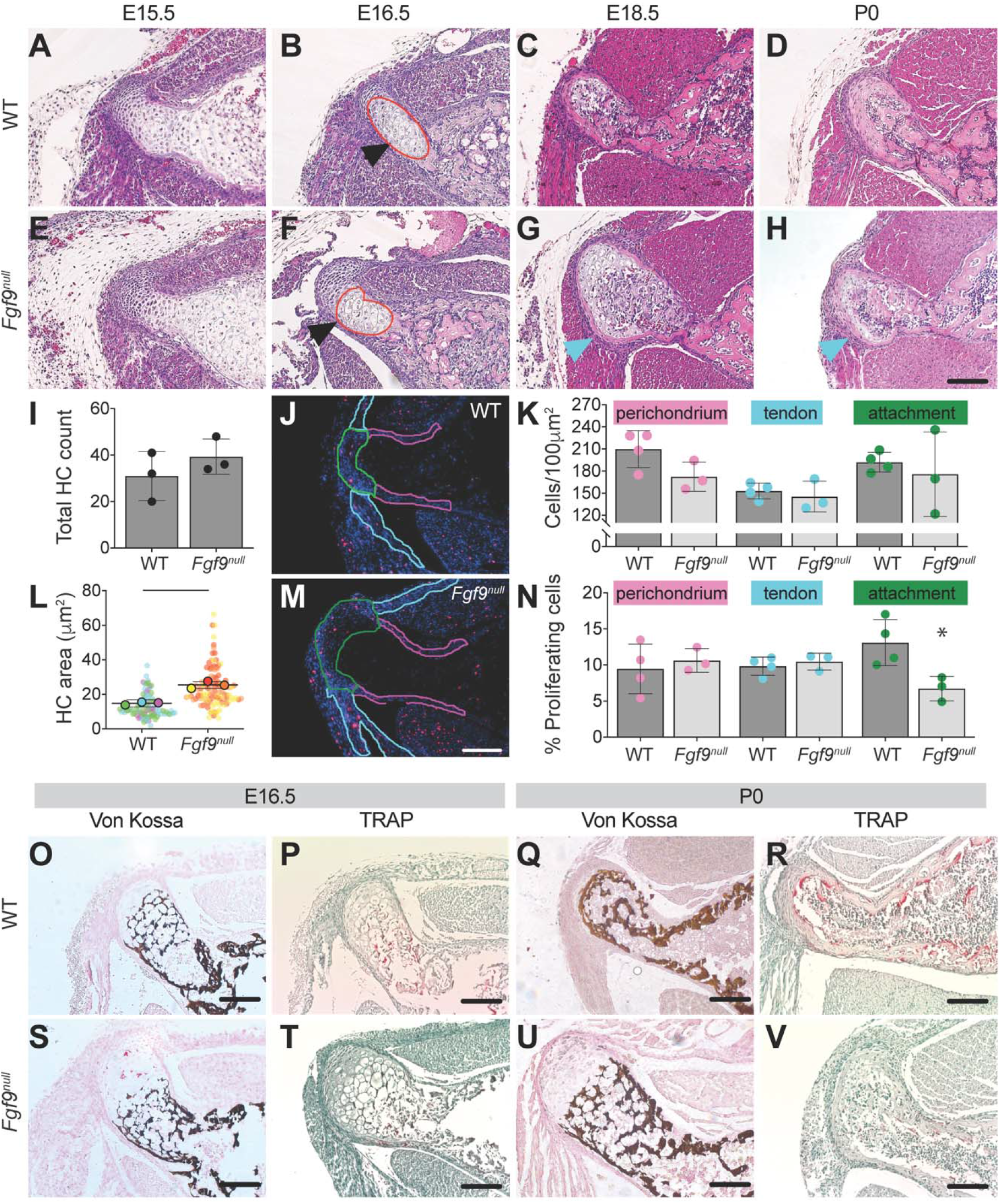
The DTs of *Fgf9^null^* embryos undergo differences in cellular and morphological development when compared to WT embryos. The DT of *Fgf9^null^* embryos was larger than the DT of WT embryos as early as E16.5 (black arrows; scale bar = 100μm). During later stages of development (E18.5 **(C, G)** and P0 **(D, H)**), hypertrophic chondrocytes (blue arrows) remained in the DT of *Fgf9^null^* embryonic limb and were absent in WT embryonic limb. H&E-stained transverse sections of the DT in **(B)** WT embryos and **(F)** *Fgf9^null^* embryos at E16.5. Red lines represent the region of hypertrophic chondrocytes. *Fgf9^null^* embryos had a **(I)** similar number of hypertrophic chondrocytes and **(L)** larger average area of hypertrophic chondrocytes compared to WT embryos. **(J,M)** Proliferating cells were labeled using EdU, in red. Outlined regions of interest include the perichondrium (pink), tendon (teal), and attachment (green). *Fgf9^null^* embryos had **(K)** similar cell density and **(N)** reduced proliferation rates at the DT attachment compared to WT embryos. **(U)** At P0 *Fgf9^null^* neonates had abnormal mineral patterning (von Kossa) and **(V)** fewer osteoclasts (TRAP) than **(Q,R)** WT neonates (scale = 100μm). Data are biological replicates (faded circles are technical replicates), mean ± SD. Lines in L and * in N indicates significant difference between groups (p < 0.05).

We visualized mineralization of the attachment using von Kossa staining and found that at E16.5, WT and *Fgf9^null^* embryos both had a woven mineral pattern within the hypertrophic region. At P0, the remodeled mineral in WT DTs formed a layer of cortical bone (Figure 2Q), whereas in *Fgf9^null^* neonates, the woven mineral pattern remained (Figure 2U). In addition, fewer osteoclasts were present at both E16.5 and P0 (Figure 2R,V; Supplemental Figure 4).

### Global loss of *Fgf9* leads to gene expression differences in muscle, but not bone

*Fgf9* is expressed as early as E10.5 in the developing limb and later in the perichondrium and muscle (Colvin et al., 1999; Garofalo et al., 1999; Hung et al., 2007; Yang and Kozin, 2009). Using FISH, we found that *Fgf9* was primarily expressed in the soft tissues surrounding the developing bone (Supplement Figure 5B). At E16.5, *Fgf9^null^* embryos had increased localized expression of *Gli, Sox9*, and *Fgf18* (markers of bone and cartilage development) within and around the DT compared to WT embryos at the same developmental stage (Figure 3B,F,N). At P0, *Fgf9^null^* neonates had reduced localized *Sox9* and *Sost* expression compared to WT neonates (Figure 3H,L).

**Figure 3:**
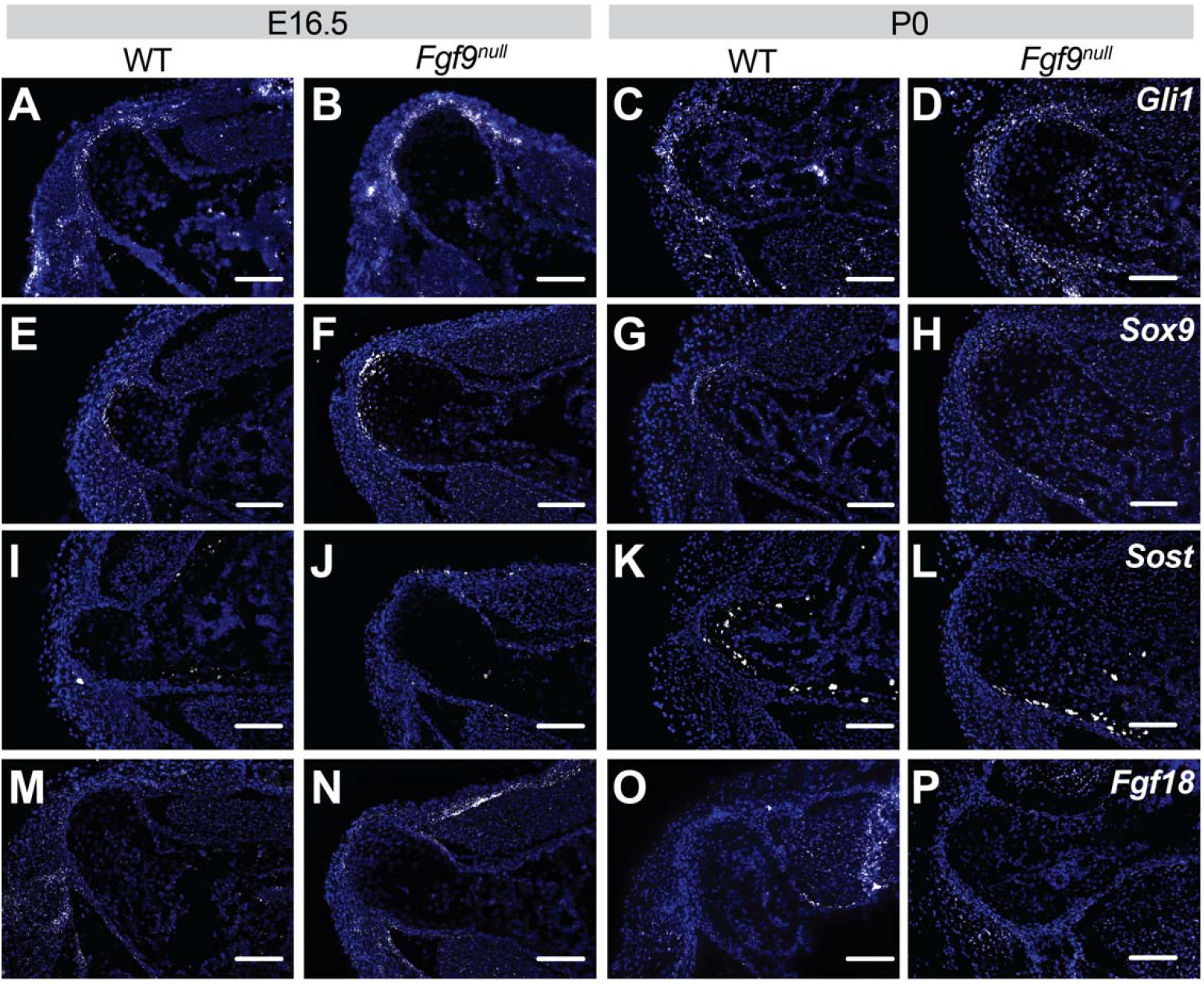
Fluorescent *in situ* hybridization (FISH) of (A-D) *Gli1*, (E-H) *Sox9*, (I-L) *Sost*, and (M-P) *Fgf18* at E16.5 and P0 of *Fgf9^null^* and WT mice. At E16.5, *Fgf9^null^* embryos had increased **(B)** *Gli,* **(F)** *Sox9*, and **(N)** *Fgf18* expression compared to **(A,E,I,M)** WT embryos. At P0, *Fgf9^null^* neonates had less **(H)** *Sox9* and **(L)** *Sost*, and similar **(D)** Gli1 and **(P)** *Fgf18* expression compared to **(C,G,L,P)** WT neonates (scale bar = 100μm).

**Figure 4:**
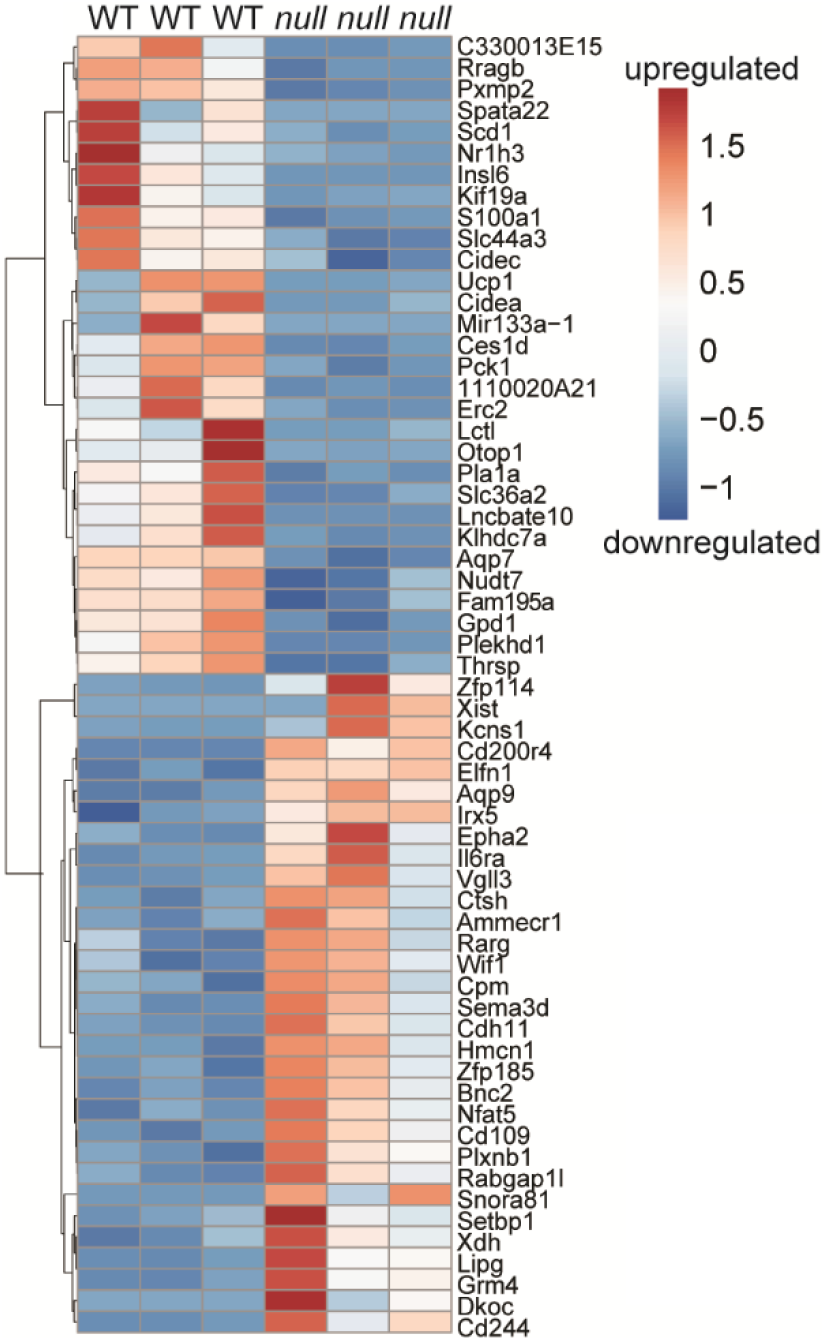
Heat map of top 30 upregulated and downregulated genes from RNA-Seq secondary analysis. 135 differentially expressed (DE) with a false discovery rate value < 0.05 and genes omitted with fold change outside 2 standard deviations of genotype average log2 fold change (FC). Color indicates z-score value as calculated and presented using Morpheus software. Scale bar = log2 FC.

For an unbiased comparison of differential gene expression from bone and muscle of WT and *Fgf9^null^* embryonic limbs, we performed bulk RNA-sequencing from micro-dissected tissue of stylopods at E18.5 (n=3/genotype). We used a cut off false discovery rate of 0.05 and discovered 805 differentially expressed (DE) genes between WT and *Fgf9^null^* muscle with only a single DE gene (*Xist*) in bone. A secondary analysis was performed to exclude DE genes with a fold change greater than two standard deviations between biological replicates. In our secondary analysis, we identified 135 unique genes (Figure 4). The primary and secondary DE gene lists were then analyzed using Database for Annotation, Visualization and Integrated Discovery (DAVID) (Huang et al., 2009a; Huang et al., 2009b) and the GOTERM cellular components direct, biological processes direct, and molecular functions direct libraries. DE genes in the primary pool of 805 genes were found in a broad set of terms (Supplemental Figure 6). DE genes in the secondary pool of 135 genes were associated with decreased expression of mitochondria/energy and lipid associated genes in *Fgf9^null^* muscle compared to WT muscle (Figure 5).

**Figure 5:**
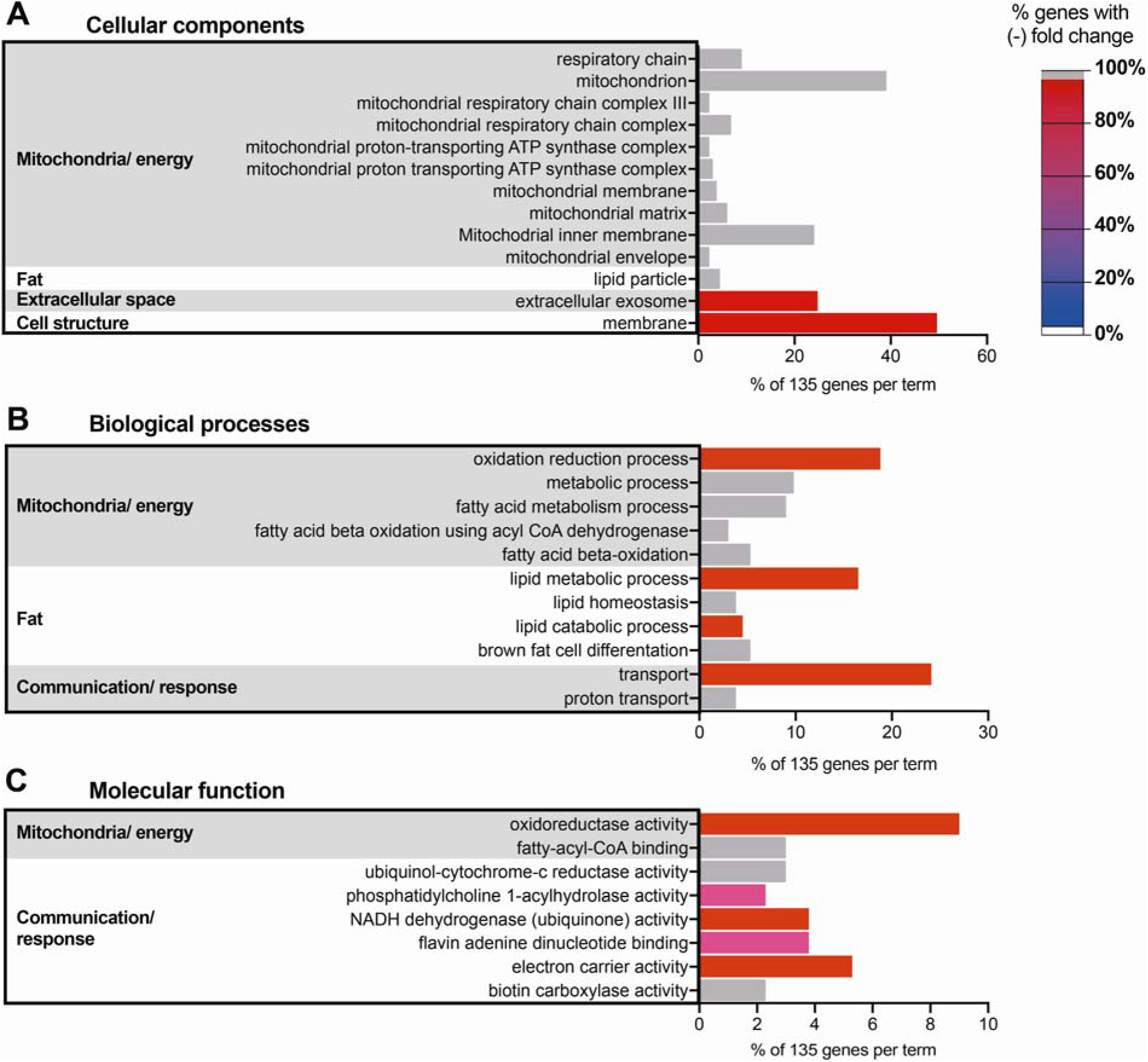
Downregulation to differentially expressed genes between *Fgf9^null^* and WT neonates related to mitochondrial and fat. 135 DE genes from secondary analyses were categorized using DAVID against a *Mus musculus* background. Terms with a Benjamini value < 0.05 were selected from the **(A)** GOTERM cellular components DIRECT, **(B)** GOTERM biological processes DIRECT, **(C)** and GOTERM molecular function DIRECT. Similar terms were then clustered into categories (bar chart = % gene in term out of 135; color = % genes in term with negative fold change). Terms with 100% negative FC (WT vs. *Fgf9^null^*) were assigned gray bars.

### Skeletal-muscle specific deletion of *Fgf9* results in enlarged tuberosities and decreased neural-related gene expression

Based on our RNAseq findings, we developed a doxycycline-inducible and skeletal-muscle specific *Fgf9^cKO^* mouse line, which, unlike *Fgf9^null^* embryos, were postnatal viable. Muscle-specific targeting was confirmed through Ai14 reporter (*Acta1*-Cre) (Figure 6GH). At 8 weeks of age, *Fgf9^cKO^* mice were similar in size and had comparable humerus lengths (Supplemental Figure 7A) compared to WT littermates. However, *Fgf9^cKO^* mice had ~35% larger DTs compared to WT littermates (Figure 6C). Bone quality of DTs between WT and *Fgf9^cKO^* mice was similar (Supplemental Figure 7B). In addition, another bony superstructure on the humerus, the lateral supracondylar ridge (LSR) near the elbow where extensor muscles originate, was ~30% larger in *Fgf9^cKO^* mice than WT mice (Figure 6D). Gene expression of *Agrn* (Figure 6I) and *Ikbkb* (Figure 6J) in deltoid muscle from *Fgf9^cKO^*mice was decreased compared to WT mice at 8 weeks but not 3 weeks of age. Acetylcholine Receptor (AChR) density (Figure 6N) was 30% significantly higher in the deltoid muscles of *Fgf9^cKO^* mice when compared to that of WT mice at 3 weeks of age (Figure 6K,L). However, AChR density normalized between genotypes by 8 weeks. There was not a significant difference in total AChR count in the deltoid for either time point (Figure 6M).

**Figure 6:**
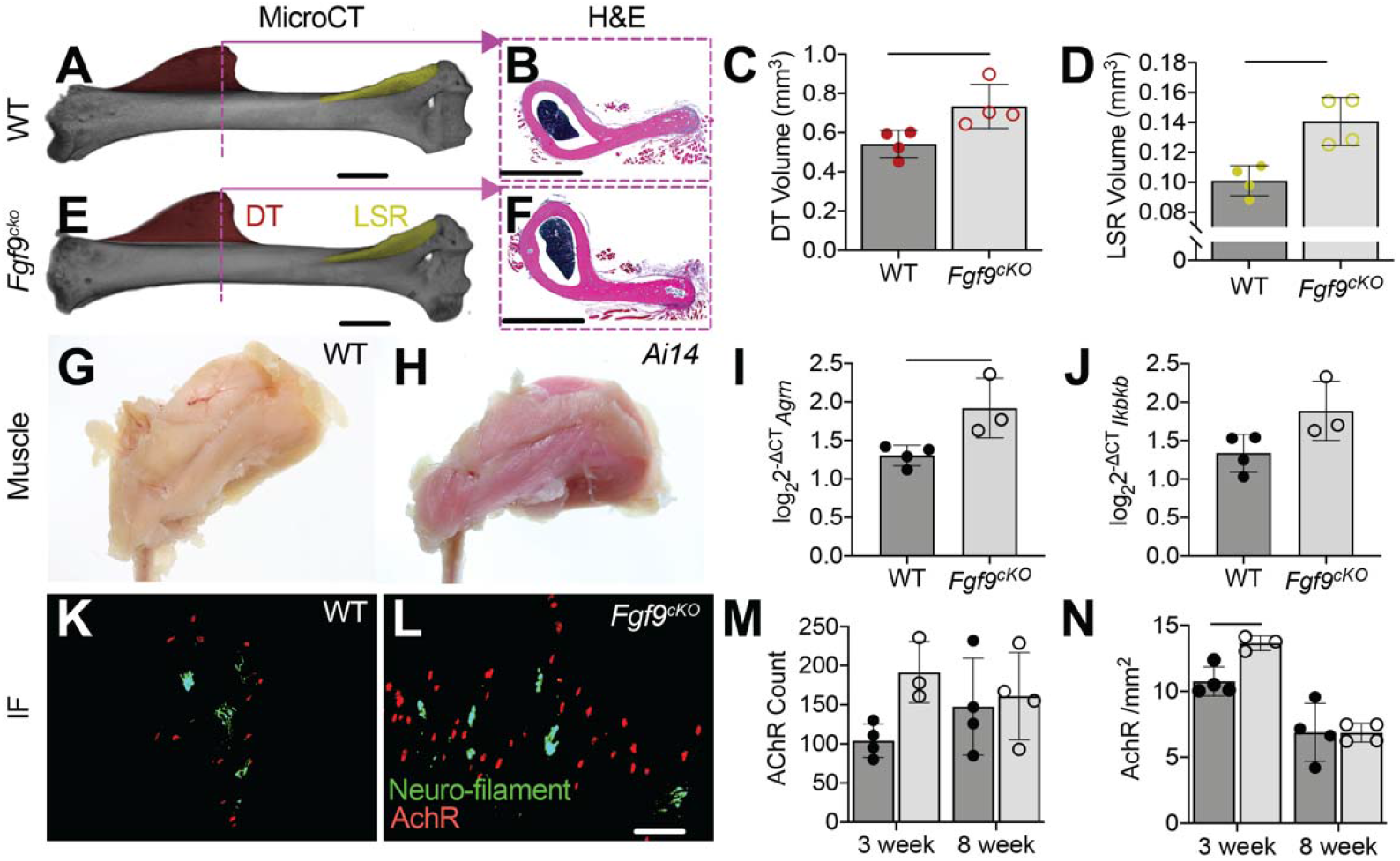
Muscle-specific knockout of *Fgf9* in mice (*Fgf9^cKO^*) resulted in larger superstructures and increased number of acetylcholine receptor (AChR) clusters in the deltoid muscle. MicroCT scans of 8 week **(A)** WT and **(E)** *Fgf9^cKO^* mice show a significantly larger DT (DT;red) and lateral supracondylar ridge (LSR; yellow) in *Fgf9^cKO^* mice. Histology of the DT support this phenotype **(B, F)**. Skeletal muscle specificity of *Acta1*-Cre was confirmed using Ai14 reporters **(H,G)**. Gene expression of **(I)** *Agrn* (p = 0.0287) and **(J)** *Ikbkb* (p = 0.0675) were increased in *Fgf9^cKO^* deltoid muscle compared to WT muscle at 8 weeks of age (ΔCT calculated using average reference gene CT of *ipo8* and *Rn18s*). Immunofluorescence (IF) was used to visualize nerves (neurofilament, green) and AChR clusters at neuromuscular junctions (α-bungarotoxin, red) in **(K)** WT and **(L)** *Fgf9^cKO^* mice. At 3 weeks postnatally, *Fgf9^cKO^* mice had significantly higher **(N)** AchR density than WT littermates and **(M)** more AChR. Data shown are biological replicates, mean ± SD. Lines in C, D, I, and N indicate significance (p < 0.05).

## Discussion

In this study, we showed that *Fgf9* may act as a negative regulator in growth of attachment superstructures such as the DT. Global deletion of *Fgf9* led to enlarged superstructure size, and these superstructures had altered endochondral ossification of the arrested growth plate. The decreased proliferation we have shown in DTs of *Fgf9^null^* embryos, as well as enlarged hypertrophic chondrocytes, is similar to findings from previous work studying long bone development of *Fgf9^null^* embryos (Hung et al., 2007; Hung et al., 2016). The prevalence of hypertrophic chondrocytes at the tendon attachment in *Fgf9^null^* embryos may be suggestive of a prolonged progenitor cell maintenance along these superstructures (Blitz et al., 2013). *Sox9* and hedgehog signaling (e.g., *Gli1*) are sequentially required for cellular expansion of growth plates during endochondral bone formation (Akiyama, 2002; Long, 2004; Mau et al., 2007; Pan et al., 2006; St-Jacques et al., 1999; Tan et al., 2018); however, migratory attachments (like the DT) form via replacement of *Sox9* expressing cells by *Gli1* expressing cells (Felsenthal et al., 2018). At E16.5, as DT enlargement begins in *Fgf9^null^* embryos, we observed increased *Gli1* and *Sox9* expression around the attachment. Indian hedgehog signaling also represses transcription of hypertrophy-related genes (Leung et al., 2011), and inhibits parathyroid hormone-like hormone (*Pthlh*) activity to maintain chondrocytes in a proliferative stage (Lanske et al., 1996). Though we saw no difference in *Pthlh* expression, the decline in cell proliferation at attachments and increased cell hypertrophy may be indicative of an acceleration in endochondral ossification. The final stage of endochondral ossification is marked by *Sost*-expressing osteocytes (Winkler et al., 2003). In our study, we observed fewer *Sost*-expressing clusters in the DT of *Fgf9^null^* neonates, suggesting that *Fgf9^null^* DTs may also maintain fewer differentiated osteocytes, which may contribute to a prolonged maintenance of chondrogenic *Gli1*+ cells at the attachment.

*Fgf9* is expressed in muscle during embryonic stages, which we and others have observed using ISH (Colvin et al., 1999; Garofalo et al., 1999; Hung et al., 2007; Yang and Kozin, 2009). Previous work has established a connection between *Fgf9* and muscle, as treatment of muscle and muscle progenitor cells with FGF9 slows maturation, enhances proliferation, and decreases expression of various myogenic genes (Huang et al., 2019). This study found supporting evidence that *Fgf9* expression in muscle may be a limiting factor in tuberosity growth. However, it remains unknown how other FGFs and their receptors, FGFRs, regulate superstructure and attachment formation.

FGF ligands have varying affinities for receptors, and receptor expression changes throughout cell development (Su et al., 2014). *Fgfr1* is highly expressed in hypertrophic chondrocytes (Peters et al., 1992), and *Fgfr2* is expressed in the condensing mesenchyme (Hall, 1987; Hall and Miyake, 2000; Roberts et al., 2019). *Fgfr1* and *Fgfr2* are co-expressed in osteogenic cells (Ornitz and Marie, 2002; Peters et al., 1992) and *Fgfr3* is predominately expressed in proliferating chondrocytes (Henderson et al., 2000). Thus, FGF signaling may play a unique and cell-specific role during establishment and growth of the *Sox9/Scx* progenitor pool found within developing superstructures (Blitz et al., 2013). Previous work in craniofacial development has shown that *Fgfr2* expression is necessary for maintaining *Sox9/Scx* progenitor pool cell fate (Roberts et al., 2019); however, spatial differences and regulatory roles of *Fgf* signaling between craniofacial and limb development are important to consider. For example, within the developing humerus, progenitor cell pools are temporally and transcriptionally different depending on location (e.g., proximal DT vs. distal olecranon superstructures) (Eyal et al., 2019). Additionally, *Fgf9* predominantly effects proximal (stylopod) skeletal elements rather than distal (zeugopod, autopod) elements (Hung et al., 2007). Thus, the timing, spatial contribution, and expression patterns of *Fgf9* on musculoskeletal development may be isolated to certain cells in proximal elements (Hung et al., 2007; Su et al., 2014).

## Material and Methods

### Mice

Prrx1-Cre, Ai14(RCL-tdT)-D (Ai14), and Acta1-rtTA-tetO-Cre (*Acta1*-Cre) transgenic mice were obtained from Jackson Laboratory (The Jackson Laboratory, Bar Harbor, ME, USA). *Fgf9*-floxed mice were generated as previously described (Colvin et al., 1999; Colvin et al., 2001). Mice carrying the *Fgf9* excised allele were maintained on a C57BL6J background. All experiments were performed with approval from the University of Delaware Institutional Animal Care and Use Committee.

### Breeding and genotyping

For global mutants, *Fgf9^ex/WT^*; *Prrx1*-Cre female mice were bred with *Fgf9^flx/flx^* male (mixed background) mice to generate pups with homozygous excised *Fgf9* alleles (*Fgf9^null^)* and *Fgf9^WT/Ex^* (WT) littermates. The exploitation of the *Prrx1*-Cre female donors resulted in global excision of floxed alleles *in utero,* thereby generating global knockouts (Colvin et al., 2001). Embryos and neonates were used for all experiments. For timed matings, we considered noon on the day following mating as 0.5 days post coitum (0.5 dpc). For muscle-specific knockouts, *Fgf9^flx/WT^*; *Acta*1-Cre mice were crossed with *Fgf9^flx/flx^* mice and dams were given doxycycline chow ad libitum throughout gestation and weaning. To confirm that muscle was being targeted, *Acta1*-Cre mice were crossed with Ai14 reporter mice and dams were given doxycycline chow ad libitum throughout gestation and weaning. Muscle targeting was confirmed through red tint to Ai14; *Acta1*-Cre mouse muscle (Figure 6H). Genotypes were determined by PCR using commercial vendors (Transnetyx, Cordova, TN, USA). The efficiency of Cre-recombination and confirmation of *Fgf9^null^* gene knockout was determined using quantitative reverse transcription-polymerase chain reaction (qRT-PCR) (WT n=3; *Fgf9^null^* n=3; Supplemental Figure 1). Samples were amplified in triplicate and averaged together to calculate and ΔCT values using *Rn18s* as the reference gene. ΔCT values were converted to log-scale (log 2^−^ΔCT). Relative expression was then calculated by dividing biological replicate values by the average WT log-scale value. Relative expression was then presented as mean ± SD. For ΔΔCT, CT values were averaged together per genotype. Per genotype, CT averages of *Fgf9*, *Rn18s*, and *Scx* were subtracted by the *Rn18s* CT to calculate the ΔCT. ΔΔCT for each gene was calculated by subtracting ΔCT for *Fgf9^null^* by ΔCT for WT. ΔΔCT was converted to fold change values (2^−^ΔΔCT), and the mean value between triplicates was used for biological replicates.

### Measurement of DT size

For global mutants, whole-mount staining was used following a standard protocol (Rigueur and Lyons, 2014). Arms of P0 neonates (WT n=7; *Fgf9^null^* n=14) were removed from whole-mount skeletons and placed in a Petri dish filled with glycerol with the anterior side facing up. In-plane with the arm, we used a ruler at the bottom of the petri dish for scaling. Samples were imaged using a Stemi dissection microscope (Carl Zeiss, Gottingen, Germany) and imaged using (EOS REBEL T5, Canon, Melville, New York, USA). The size of the DT (Supplemental Figure 2A) and humerus (Supplemental Figure 2C) were measured using ImageJ (Schneider et al., 2012). When we observed no significant difference in mineralized humerus area and DT area combined (Supplemental Figure 2B), we based DT size on the ratio of DT area to mineralized humerus area. Red mineral staining determined the mineralized zone, and DT measurement included any cartilage observed at the crest of the DT. We determined the relative placement of the DT based on a ratio of length distal to DT over the length of the humerus. Both mineralized humerus and full humerus were measured. The distal length was based on the distance between the most distal part of the DT and the most distal part of the mineralized/full humerus. Humerus length was based on the distance between the most proximal part of the mineralized/full humerus and the most distal part of the mineralized/full humerus. Once we determined no difference between left and right arms, values were averaged together.

### Histology and *in situ* hybridization

Embryonic and neonatal mice were eviscerated and fixed in 4% paraformaldehyde (PFA) for 24 hours. Forelimbs for histology of a subset of embryonic age 18.5 (E18.5) embryos and P0 neonates were decalcified using formic acid (Formical 2000, Statlab, McKinney, TX, USA) for 24 hours. Histomorphometry was carried out on 7 μm paraffin sections of samples from E15.5 to P0 (n= at least 3 per genotype/time point). Samples were stained using hematoxylin and eosin (H&E), coverslipped with acrymount (Statlab, McKinney, TX, USA), and imaged with an Upright microscope (Carl Zeiss, Gottingen, Germany). The area and number of hypertrophic chondrocytes were measured in the DT of E16.5 embryos using the freehand and multipoint tools in ImageJ (Schneider et al., 2012). Non-decalcified E16.5 and P0 samples had mineralized bone visualized by von Kossa staining (n=3 per age and genotype) with Nuclear Fast Red counterstaining as part of an established protocol (Abcam, Cambridge, UK). For osteoclast staining, P0 neonates were decalcified using 14% ethylene-diamine-tetraacetic acid (EDTA), paraffin sectioned, and stained for TRAP (n=3 per age and genotype). We performed *in situ* hybridization using the RNAscope Multiplex Fluorescent Reagent Kit v2 (Advanced Cell Diagnostics, Hayward, CA, USA). Non decalcified E16.5 and P0 samples (n=3 per age and genotype) were treated with *Fgf9, Fgf18, Gli1, Pthlh, Scx, Sost, Sox9, and Wnt7b* probes (Advanced Cell Diagnostics, Hayward, CA, USA) for 2 hours at 40C. A positive control slide used a mixture of probes targeting housekeeping genes *Polr2A, Ppib,* and *Ubc*. A negative control slide used dapB (a bacteria gene). Nuclei were counterstained with DAPI and mounted with Citifluor Antifade mounting medium (Electron Microscopy Science, Hatfield, PA, USA). Slides were imaged using Zeiss Axio-observer Z1 microscope (Axio.Observer Z1, Carl Zeiss, Gottingen, Germany).

### Immunohistochemistry

Embryonic and neonatal mice were eviscerated and fixed in 4% PFA for 24 hours (n=3 age and genotype). Forelimbs for histology of P0 neonates were decalcified using 14% EDTA for 24 hours. Histomorphometry was carried out on 7 μm paraffin sections of samples from E15.5 and P0 (n= at least 3 per genotype/time point). Slides were deparaffinized and hydrated to 70% ETOH, followed by heat mediated antigen retrieval (Sodium Citrate Buffer, pH 6.0) at 65°C for 2 hours. Slides were kept in the solution until equilibrated to room temperature, quenched in 0.3% hydrogen peroxide (Santa Cruz, Dallas, TX) for 30 minutes and blocked with 5% goat serum in 1% phosphate buffer saline in room temperature for 1 hour. For primary antibody, rabbit monoclonal anti-collagen X antibody (ab260040; 1:1000 dilution) (ABCAM, Cambridge, MA, USA) was used for overnight incubation at 4°C. Following incubation, HRP/DAB system (Millipore Sigma) was used for detection. Samples were then counterstained with hematoxylin and mounted with a xylene-based mounting medium (Acrymount, Stat Lab, McKinney, TX, USA) and imaged using a Lionheart FX (BioTek, Winooski, Vermont, USA). 2 P0 (n=1 per genotype) samples were omitted from analysis due to surplus nonspecific staining.

### EdU proliferation

A dam at E16.5 in gestation was given an intraperitoneal (IP) injection with EdU (5-ethynyl-2’-deoxyuridine) from the Click-iT^TM^ EdU Alexa FluorTM 647 Imaging Kit (Invitrogen, Carlsbad, CA, USA) at a dosage of 0.25mg /10g body weight. The dam was euthanized two hours later through CO2 asphyxiation. Embryos were eviscerated and fixed in 4% PFA for 24 hours. Forelimbs were processed for paraffin histology, and humeri were cut in the transverse plane at 7 μm. Proliferating cells were stained with Alexa Fluor 647 and Hoechst 33342 as a counterstain. Slices were mounted with Citifluor Antifade Mountant. Two histological sections per sample treated with EdU (WT n=4; *Fgf9^null^* n=3) were imaged using a Zeiss Axio-observer Z1 microscope (Axio.Observer. Z1, Carl Zeiss, Gottingen, Germany). Differential interference contrast (DIC) images and fluorescence images were collected from the same observed tissue to visualize tissue morphology. DIC allowed us to observe the highly organized structure of the tendon and perichondrium. Regions were separated into perichondrium, tendon, and attachment. Tissue area, nuclei number, and the number of EdU positive proliferating cells were measured and counted within ImageJ using the freehand and multipoint tools. Two slides per sample were averaged together. For the tendon and perichondrium, tissues on the medial and lateral side of the DT were measured separately. Once no phenotypic difference based on the side of tissue was observed, medial and lateral values were summed together.

### Evaluation of bone deposition and resorption

For dynamic bone histomorphometry, Calcein (Sigma, C0875; 15 mg/kg body weight; 12 pm) was injected into pregnant females intraperitoneally (IP) at E16.5 and Alizarin Complexone (Sigma, a3882; 45 mg/kg IP; 2 days later at 6 pm) at E18.5 with a set time in between of 42 hours. Neonates were collected at birth, eviscerated, and fixed for 24 hours in 4% PFA. After paraffin embedding, samples were sectioned at 7um. Slides were deparaffinized, dehydrated, and mounted with Acrymount. Slides were imaged using Zeiss Axio-observer Z1 microscope (Axio.Observer. Z1, Carl Zeiss, Gottingen, Germany). Calcein Green and Alizarin Red were imaged using fluorescence microscopy and quantified using ImageJ by using the freehand tool to measure the area of green and red stained mineral. The mineralization area (MA) of the DT was measured, and the mineral growth rate was calculated using:

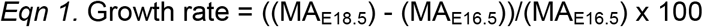

### Bulk RNA-Sequencing

E18.5 embryos were collected, toed, and eviscerated under sterile conditions. RNA was extracted from each sample (n=3 WT and n = 3 *Fgf9^null^*) using Trizol/chloroform and spin-column assembly with on-column DNase treatment (RNeasy mini kits; Qiagen Germantown, MD). We assessed the quality of RNA using fragment analysis to obtain RNA integrity number (RIN) before proceeding with Library preparation (NEBNext® Ultra™ RNA Library Prep Kit, Illumina, San Diego, CA, USA). Each sample was sequenced in Illumina HiSeq 2000 (Single read, 50 base pair). Sequence reads from each sample were quality assessed (FastQC) & all the reads were quality trimmed and filtered (threshold length of the sequence was 30 & Phred score of each base was 28). Reads were mapped to the *Mus musculus* reference genome (mm10) using TopHat2 with default settings (ver 2.1.0)(Kim et al., 2013) and mapping quality control metrics were obtained fro RSeQC (ver 2.6.1)(Wang et al., 2012). Per gene counts were determined using HTseq (ver 0.6.1p1)(Anders et al., 2015) and used for pairwise determination of differential expression analyses using EdgeR (ver 3.28.1) (Robinson et al., 2010) with a significance cutoff of FDR-corrected p<0.05. To control for consistency within biological replicates, a secondary analysis was performed to exclude DE within a group if the mean count per million for a gene feature in a given sample was >log2FC different than its biological replicates (yielding 135 genes). These 135 genes were then uploaded to Morpheus, which assigned a z score to determine the top 30 upregulated and downregulated genes and generate a heatmap (Morpheus, https://software.broadinstitute.org/morpheus). Both sample pools (primary analysis of 805 and secondary analysis of 135) were then uploaded to DAVID Bioinformatics Resources 6.8 (Huang et al., 2009a; Huang et al., 2009b) and run against a mm10 background. GOTERM cellular components direct, GOTERM biological processes direct, and GOTERM molecular functions direct libraries were examined, and terms were ranked based on Benjamini number (DAVID adjusted p value) (Benjamini and Hochberg, 1995). Terms with a Benjamini number < 0.05 were percentiled and categorized based on similarity.

### Skeletal Muscle Immunofluorescence

Muscle-specific *Fgf9* knockouts and WT littermates were euthanized at 3 weeks of age (n=4 WT, n=3 *Fgf9^cKO^*) and 8 weeks of age (n=4 WT, n=4 *Fgf9^cKO^*) for histology, structural analysis, and RNA isolation. Immediately following euthanasia, the right deltoid muscles of each mouse were dissected in an RNase-free environment, flash-frozen in liquid nitrogen, and stored at −80°C until RNA isolation. The left deltoid muscles were then dissected, fixed in 4% PFA for 20 minutes, cryo-protected, and then embedded in optimal cutting temperature (OCT) media for cryosectioning (longitudinal plane). The left humerus from each mouse was also dissected, fixed in 4% PFA, and stored in 70% ethanol.

The immunofluorescence staining protocol used in this study was based on previously established protocol (Mead et al., 2011). For 3-week and 8-week old deltoids, 35 μm thick sections were used immunohistochemistry (IHC) of acetylcholine receptors (α-Bungarotoxin extracted from Bungarus muticinctus, conjugated to Alexa Fluor 647, Thermofisher #B35450) and nerve fibers (chicken anti-neurofilament-H antibody, Thermofisher #PA1-10002; Alexa Fluor 488 donkey anti-chicken, Jackson Immunofluorescence #703-545-155). Samples were then imaged using a Zeiss Axio-observer Z1 microscope (Axio.Observer Z1, Carl Zeiss, Gottingen, Germany) at 20X magnification subfields which were then stitched in order to obtain a complete image of the ventral, middle, and dorsal heads of the deltoid. Images were visualized using fluorescent microscopy (neurofilament conjugated to 488 nm and neuromuscular junctions conjugated to 647 nm). AChR were counted in ImageJ (Schneider et al., 2012) using the multipoint tool. Muscle area was collected by opening the file in Dragonfly software (Object Research Systems, Montreal, Quebec, Canada) and separated by color channel. Muscle regions of interest were selected and area measured based on autofluorescence in GFP channel.

### Micro-computed Tomography

Micro-computed tomography (microCT) was used to obtain 3-dimensional renderings of the left humerus of 8-week old mice (n=4 WT, n=4 *Fgf9^cKO^*) to visualize topographical data on DT volume, LSR volume, DT bone volume/ total volume (BV/TV), humerus diaphysis length. MicroCT scans were performed using a Scanco μCT 35 (X-ray energy: 60kV; X-ray intensity: 95ma; integration time: 1100msec; pixel size: 10.6mm; rotation step: 0.4°)(Bouxsein et al., 2010). All raw scans were reconstructed and analyzed using Dragonfly software (Object Research Systems, Montreal, Quebec, Canada). Scans were aligned down the humeral axis and a box form was created to extract the bone ridges of interest. Using the spit at OTSU function, bone volume was selected based on pixel value. The bone analysis function was then used to fill in the bone volume and give the total volume. Diaphysis length was measured from the proximal growth plate to the Coronoid Fossa. After scans were obtained, humeri were decalcified in 14% EDTA, processed for paraffin embedding, sectioned at 7um thick, and stained with H&E. One *Fgf9^cKO^* sample and one WT sample were omitted due to processing errors.

### Neural/Muscle-related Gene Expression

*Fgf9^cKO^* and WT littermates were aged to 3 weeks (n=3 WT, n=3 *Fgf9^cKO^*) and 8 weeks of age (n=4 WT, n=3 *Fgf9^cKO^*). qRT-PCR was used to compare differential gene expression of skeletal muscle using muscle contractility genes (*Dmpk, Mef2c*, and *Ikbkb*) and neural genes (*Nes, Agrn, Gap43, Ache, Rapsn, Chrne*, and *Musk*). Neural genes were selected for their role in the development and stabilization of neurons and neuromuscular junctions as well as for their role in the signal propagation and firing pathway. Contractility genes were chosen for their role in encoding genes necessary for cell-cell communication and differentiation, skeletal muscle contraction and relaxation, myogenesis, and homeostasis (Bakkar et al., 2008). qRT-PCR, samples were amplified in triplicate and averaged together to calculate and ΔCT values with the average ΔCT of *ipo8* and *Rn18s* serving as the reference gene value. ΔCT values were converted to log-scale (log2ΔCT). Log scale was then presented as mean ± SD.

### Statistical Analysis

Statistical analyses were performed in Prism (Prism GraphPad 6.0, LaJolla, CA, USA). For whole-mount area measurements, DT area was divided by the humerus area. For relative DT placement, distal length was divided by the whole length across the mineralized and whole humerus. The resulting ratios were compared through separate two-sample t-tests. Hypertrophic cell area measurements were averaged together per mouse and a two-sample t-test compared the number of hypertrophic chondrocytes and the average hypertrophic chondrocyte area between *Fgf9^null^* embryos and WT embryos. A two-way ANOVA was run to compare cell density, proliferating cell density, and percent of proliferating cells for the EdU proliferation assay. The two-way ANOVA looked for differences between WT and *Fgf9^null^* embryos in the perichondrium, tendon, and deltoid-tendon attachment. Whole-mount, hypertrophic cell, and EdU quantifications were repeated by a separate observer through coded files to minimize bias. Two-way ANOVA was used to compare the osteoclast number between WT and *Fgf9^null^* samples at E16.5 and P0. For dynamic bone histomorphometry, 3 slides per sample were measured, and mineral growth rate was calculated. Technical replicates were averaged together per sample, and a two-sample t-test was performed. A two-sample t-test was used to compare *Fgf9^cKO^* gene expression for supplemental figures 8 and 9. A two-sample t-test was used to analyze LSR volume, DT volume, DT BV/TV, and humeri length between *Fgf9^cKO^* and WT mice at 8 weeks. Two-way ANOVA was used to compare gene expression and AchR clustering between *Fgf9^cKO^* and WT mice at 3 week and 8-week time points.

## Acknowledgments

We thank Brewster Kingham, Summer Thompson, and Mark Shaw in the University of Delaware DNA Sequencing & Genotyping Center for assistance with for assistance with RNA sequencing; Shaneaka Anderson, Kierstyn Hendricks, and Kayleigh Denner for help with histology; Crystal Idelburg (Washington University) for TRAP staining; Karol Miaskiewicz for high performance computing support.

## Supplemental Figures

**Supplemental Figure 1:**
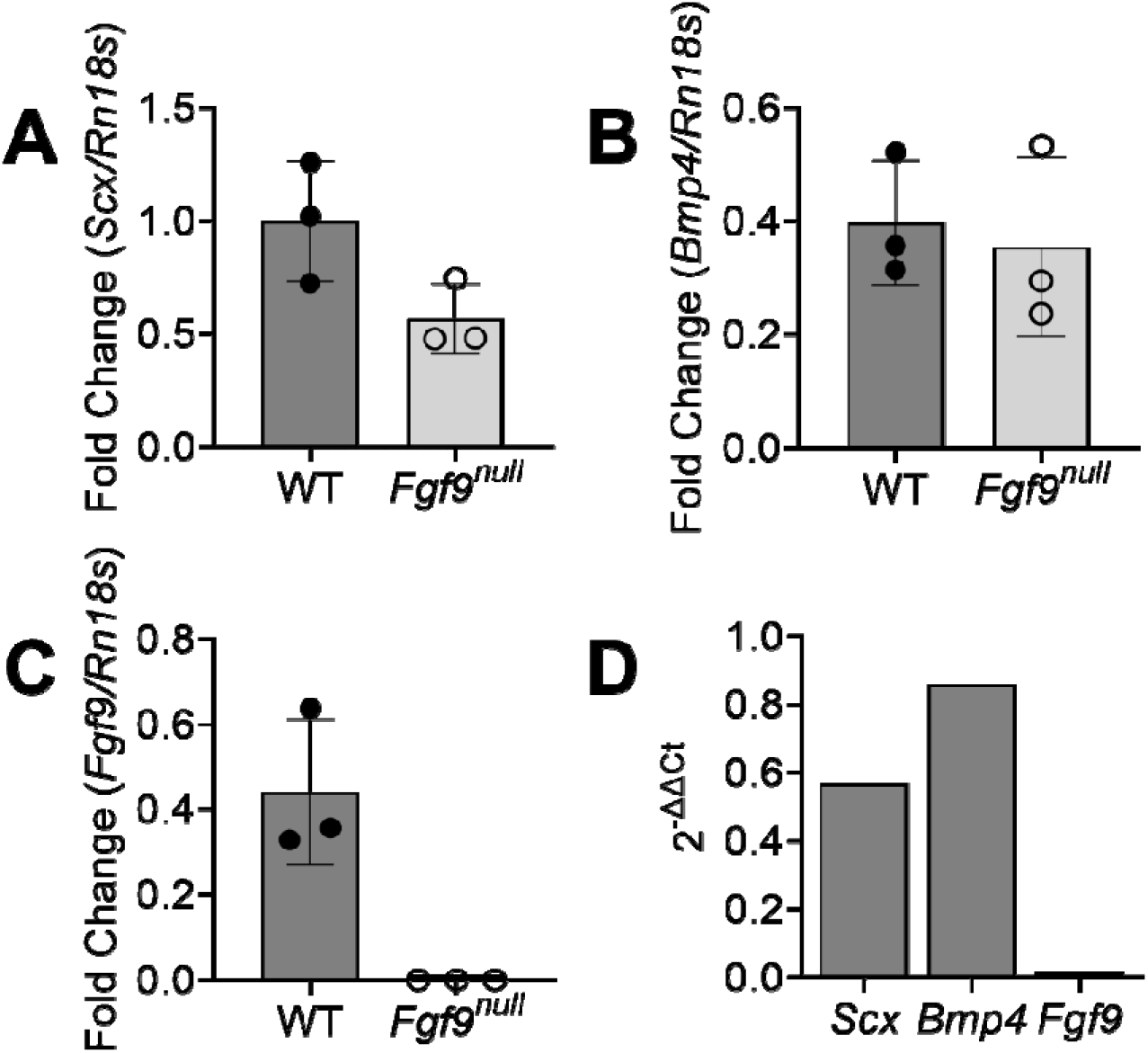
No *Fgf9* expression in *Fgf9^null^* E18.5 embryos when compared to WT E18.5 embryos. qRT-PCR analyses show embryonic *Fgf9^null^* muscle had (A) reduced *Scx* expression, **(B)** similar *Bmp4* expression, and **(C,D)** no expression of *Fgf9* when compared to WT embryonic muscle. Fold change are biological replicates (2^-^ΔCT), mean ± SD. 2^−^ΔΔCT are mean (ΔΔCT= *Fgf9^null^* (gene of interest CT averages -*Rn18s* CT average) - WT (gene of interest CT averages - *Rn18s* CT average)).

**Supplemental Figure 2:**
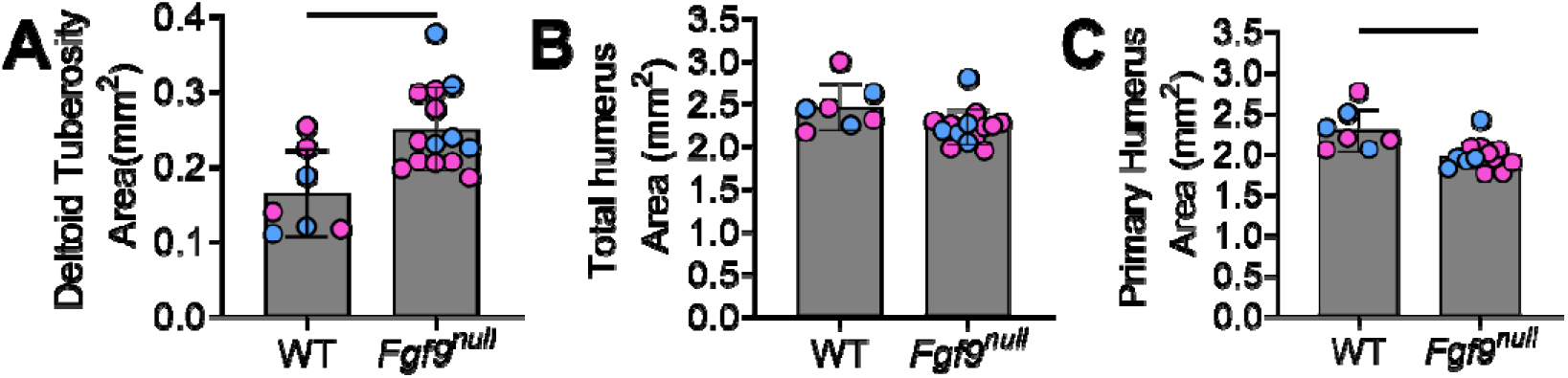
*Fgf9^null^* have a larger DT area with smaller primary humerus area than WT neonates. **(A)** Area of the DT was significantly larger in *Fgf9^null^* neonates compared to WT neonates at P0. Whether the mice were male (blue) or female (pink) had no effect. **(B)** There was no difference in the combined area of the DT and primary humerus between *Fgf9^null^* and WT neonates. **(C)** The area of just the primary humerus was significantly smaller in *Fgf9^null^* neonates compared to WT neonates. Data are biological replicates, mean ± SD. Lines in A and C indicate significance (p < 0.05).

**Supplemental Figure 3:**
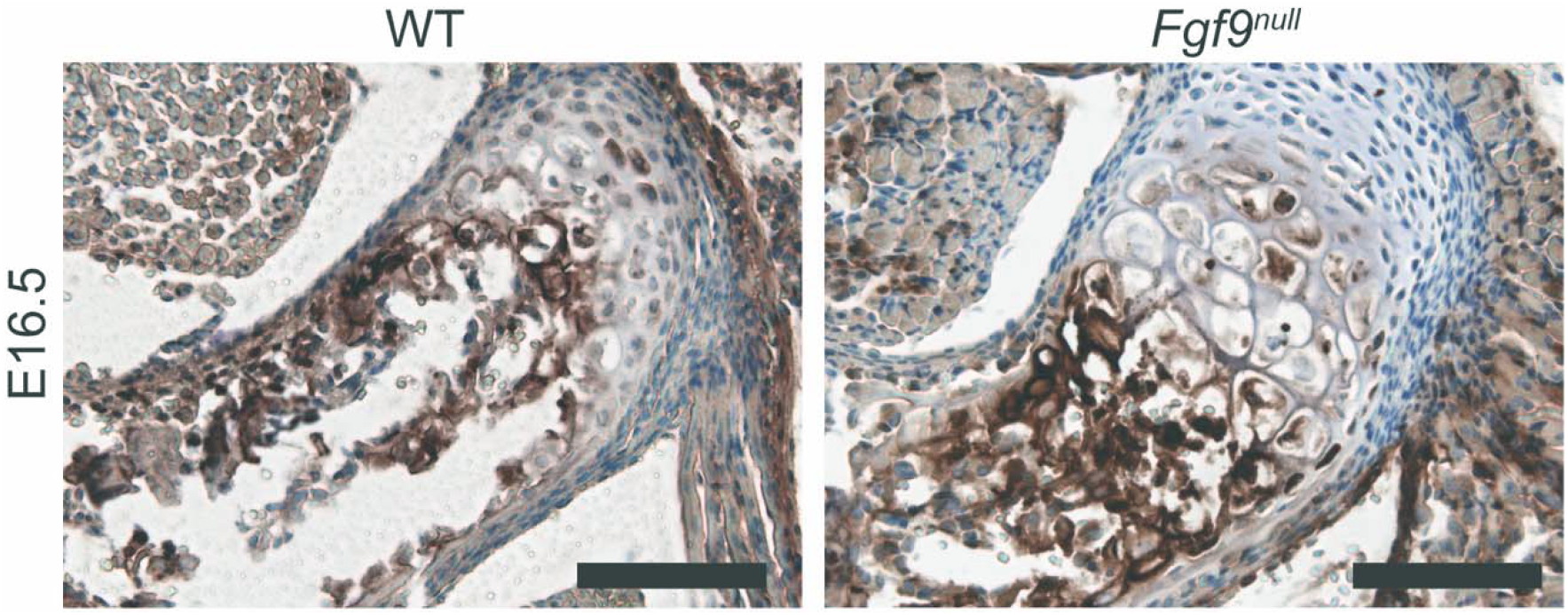
Type X collagen immunostaining of the DT in WT and *Fgf9^null^* embryos at E16.5. The tendon attachment has less ColX staining but more ColX+ hypertrophic cells in *Fgf9^null^* attachments compared to WT attachments (scale bar = 100μm).

**Supplemental Figure 4:**
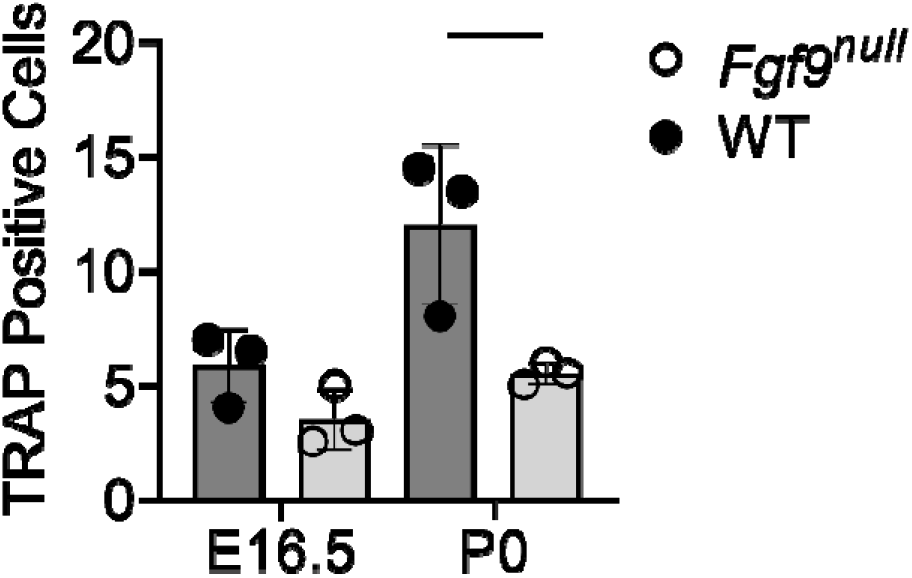
Fewer osteoclasts present in *Fgf9^null^* DTs. *Fgf9^null^* mice have fewer osteoclasts at E16.5 and significantly fewer osteoclasts at p0. Data are biological replicates (1-2 technical replicates averaged per sample) mean ± SD. Line indicate significance (p < 0.05).

**Supplemental Figure 5:**
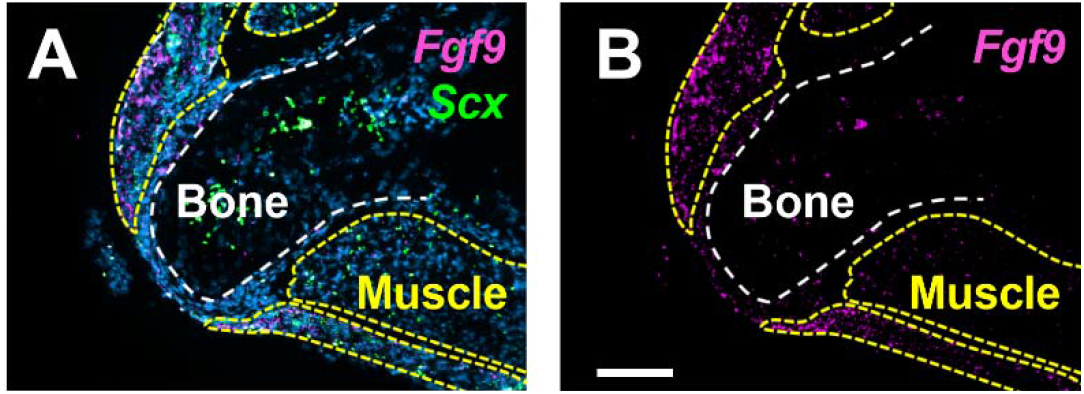
Multiplexed FISH shows that *Fgf9* is primarily expressed in the surrounding soft tissues, rather than bone. **(A,B)** *Fgf9* was observed mostly in the surrounding skeletal muscle when compared to the inside of the DT (scale bar = 100μm).

**Supplemental Figure 6:**
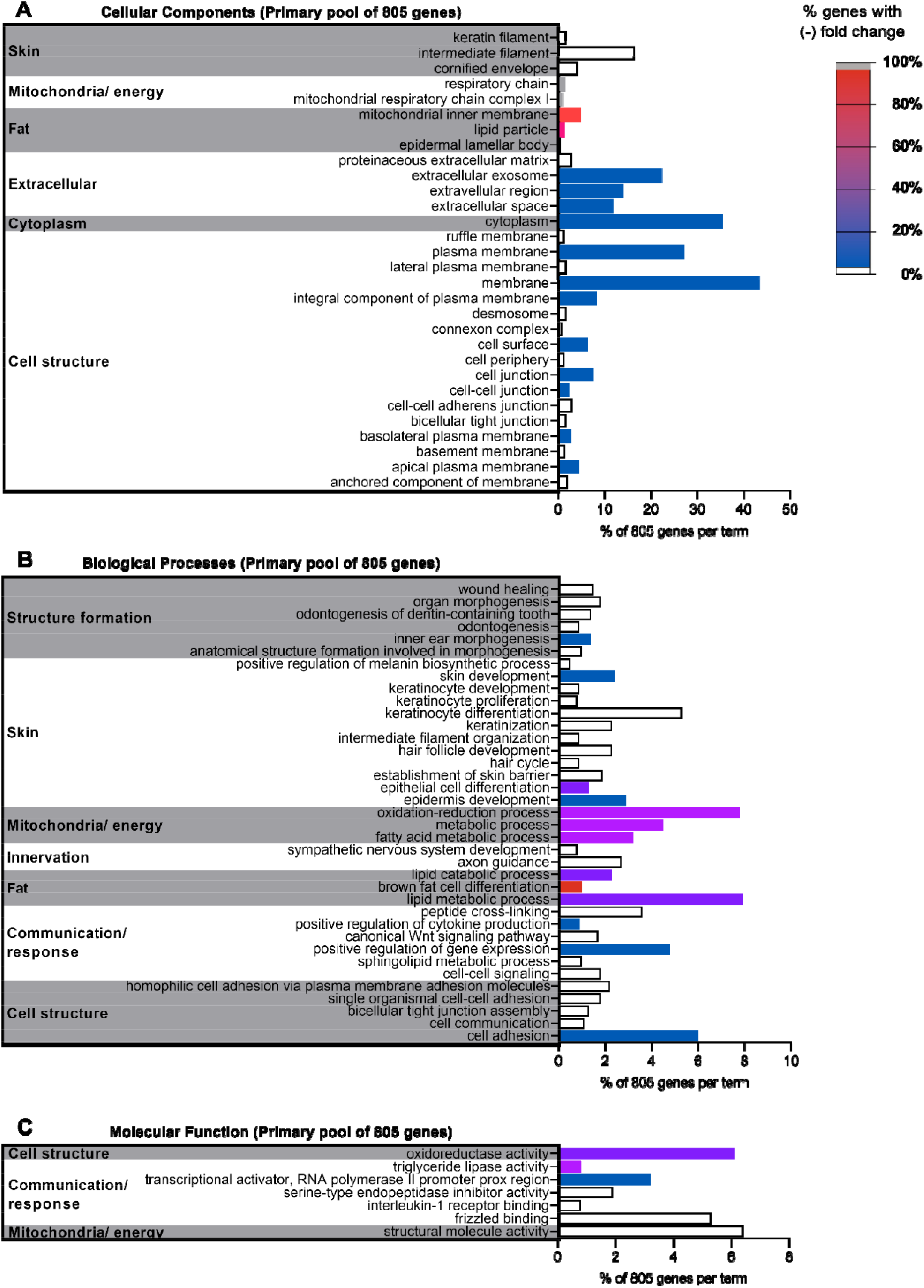
DAVID analysis of primary pool of DE genes between *Fgf9^null^* and WT neonates yield diverse terms. 805 DE genes were uploaded into DAVID against a *Mus musculus* background. Terms with a Benjamini value < 0.05 were selected from the **(A)** GOTERM cellular components DIRECT, **(B)** GOTERM biological processes DIRECT, **(C)** and GOTERM molecular function DIRECT. Similar terms are then clustered into categories (Bar chart = % gene in term out of 805; color = % genes in term with negative FC).

**Supplemental Figure 7:**
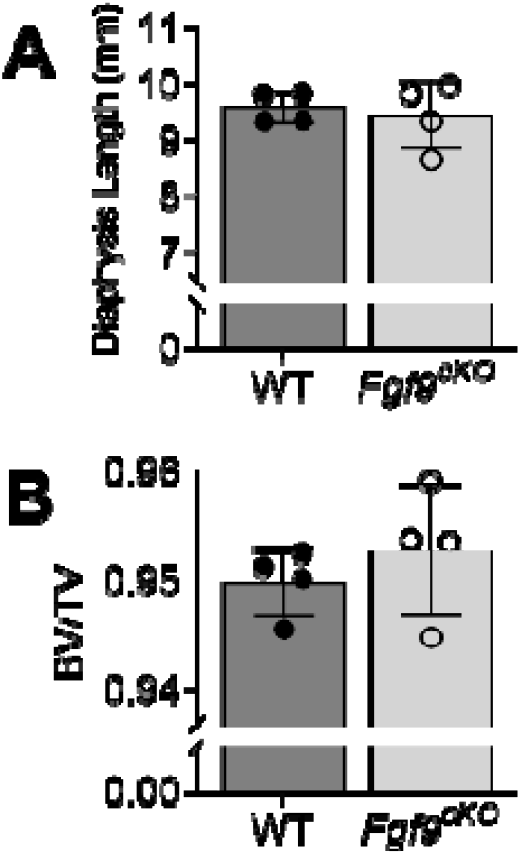
No difference in humerus length and DT Bone volume/Total volume (BV/TV) between *Fgf9^cKO^* and WT mice. From MicroCT scans of 8 week samples, **(A)** *Fgf9^cKO^* mice had a humerus of similar length when compared to WT littermates. **(B)** Though the DT was larger, BV/TV in *Fgf9^cKO^* and WT littermates. Data are biological replicates, mean ± SD.

## Works Cited

Akiyama, H. (2002). The transcription factor Sox9 has essential roles in successive steps of the chondrocyte differentiation pathway and is required for expression of Sox5 and Sox6. Genes & Development 16, 2813–2828.

Anders, S., Pyl, P. T. and Huber, W. (2015). HTSeq--a Python framework to work with high-throughput sequencing data. Bioinformatics 31, 166–169.

Arvind, V. and Huang, A. H. (2017). Mechanobiology of limb musculoskeletal development. Ann. N. Y. Acad. Sci. 1409, 18–32.

Bakkar, N., Wang, J., Ladner, K. J., Wang, H., Dahlman, J. M., Carathers, M., Acharyya, S., Rudnicki, M. A., Hollenbach, A. D. and Guttridge, D. C. (2008). IKK/NF-κB regulates skeletal myogenesis via a signaling switch to inhibit differentiation and promote mitochondrial biogenesis. Journal of Cell Biology 180, 787–802.

Benjamin, M., Toumi, H., Ralphs, J. R., Bydder, G., Best, T. M. and Milz, S. (2006). Where tendons and ligaments meet bone: attachment sites (‘entheses’) in relation to exercise and/or mechanical load. Journal of Anatomy 208, 471–490.

Benjamini, Y. and Hochberg, Y. (1995). Controlling the False Discovery Rate: A Practical and Powerful Approach to Multiple Testing. Journal of the Royal Statistical Society: Series B (Methodological) 57, 289–300.

Berendsen, A. D. and Olsen, B. R. (2015). Bone development. Bone 80, 14–18.

Biewener, A. A., Fazzalari, N. L., Konieczynski, D. D. and Baudinette, R. V. (1996). Adaptive changes in trabecular architecture in relation to functional strain patterns and disuse. Bone 19, 1–8.

Blitz, E., Viukov, S., Sharir, A., Shwartz, Y., Galloway, J. L., Price, B. A., Johnson, R. L., Tabin, C. J., Schweitzer, R. and Zelzer, E. (2009). Bone ridge patterning during musculoskeletal assembly is mediated through SCX regulation of Bmp4 at the tendon-skeleton junction. Dev Cell 17, 861–873.

Blitz, E., Sharir, A., Akiyama, H. and Zelzer, E. (2013). Tendon-bone attachment unit is formed modularly by a distinct pool of Scx- and Sox9-positive progenitors. Development 140, 2680–2690.

Bouxsein, M. L., Boyd, S. K., Christiansen, B. A., Guldberg, R. E., Jepsen, K. J. and Müller, R. (2010). Guidelines for assessment of bone microstructure in rodents using micro-computed tomography. J Bone Miner Res 25, 1468–1486.

Colvin, J. S., Feldman, B., Nadeau, J. H., Goldfarb, M. and Ornitz, D. M. (1999). Genomic organization and embryonic expression of the mouse fibroblast growth factor 9 gene. Dev. Dyn. 216, 72–88.

Colvin, J. S., White, A. C., Pratt, S. J. and Ornitz, D. M. (2001). Lung hypoplasia and neonatal death in Fgf9-null mice identify this gene as an essential regulator of lung mesenchyme. Development 128, 2095–2106.

Coutu, D. L., François, M. and Galipeau, J. (2011). Inhibition of cellular senescence by developmentally regulated FGF receptors in mesenchymal stem cells. Blood 117, 6801–6812.

Deng, C., Wynshaw-Boris, A., Zhou, F., Kuo, A. and Leder, P. (1996). Fibroblast growth factor receptor 3 is a negative regulator of bone growth. Cell 84, 911–921.

Eyal, S., Kult, S., Rubin, S., Krief, S., Felsenthal, N., Pineault, K. M., Leshkowitz, D., Salame, T.-M., Addadi, Y., Wellik, D. M., et al. (2019). Bone morphology is regulated modularly by global and regional genetic programs. Development 146, dev167882.

Felsenthal, N., Rubin, S., Stern, T., Krief, S., Pal, D., Pryce, B. A., Schweitzer, R. and Zelzer, E. (2018). Development of migrating tendon-bone attachments involves replacement of progenitor populations. Development 145,.

Garofalo, S., Kliger-Spatz, M., Cooke, J. L., Wolstin, O., Lunstrum, G. P., Moshkovitz, S. M., Horton, W. A. and Yayon, A. (1999). Skeletal dysplasia and defective chondrocyte differentiation by targeted overexpression of fibroblast growth factor 9 in transgenic mice. J. Bone Miner. Res. 14, 1909–1915.

Govindarajan, V. and Overbeek, P. A. (2006). FGF9 can induce endochondral ossification in cranial mesenchyme. BMC Dev. Biol. 6, 7.

Hall, B. K. (1987). Earliest evidence of cartilage and bone development in embryonic life. Clin. Orthop. Relat. Res. 255–272.

Hall, B. K. and Miyake, T. (2000). All for one and one for all: condensations and the initiation of skeletal development. Bioessays 22, 138–147.

Hamburger, V. and Waugh, M. (1940). The Primary Development of the Skeleton in Nerveless and Poorly Innervated Limb Transplants of Chick Embryos. Physiological Zoology 13, 367–382.

Hamrick, M. W., McPherron, A. C. and Lovejoy, C. O. (2002). Bone Mineral Content and Density in the Humerus of Adult Myostatin-Deficient Mice. Calcif Tissue Int 71, 63–68.

Henderson, J. E., Naski, M. C., Aarts, M. M., Wang, D., Cheng, L., Goltzman, D. and Ornitz, D. M. (2000). Expression of FGFR3 with the G380R Achondroplasia Mutation Inhibits Proliferation and Maturation of CFK2 Chondrocytic Cells. J Bone Miner Res 15, 155–165.

Huang, D. W., Sherman, B. T. and Lempicki, R. A. (2009a). Bioinformatics enrichment tools: paths toward the comprehensive functional analysis of large gene lists. Nucleic Acids Research 37, 1–13.

Huang, D. W., Sherman, B. T. and Lempicki, R. A. (2009b). Systematic and integrative analysis of large gene lists using DAVID bioinformatics resources. Nat Protoc 4, 44–57.

Huang, J., Wang, K., Shiflett, L. A., Brotto, L., Bonewald, L. F., Wacker, M. J., Dallas, S. L. and Brotto, M. (2019). Fibroblast growth factor 9 (FGF9) inhibits myogenic differentiation of C2C12 and human muscle cells. Cell Cycle 18, 3562–3580.

Hung, I. H., Yu, H., Lavine, K. J. and Ornitz, D. M. (2007). FGF9 regulates early hypertrophic chondrocyte differentiation and skeletal vascularization in the developing stylopod. Dev Biol 307, 300–313.

Hung, I. H., Schoenwolf, G. C., Lewandoski, M. and Ornitz, D. M. (2016). A combined series of Fgf9 and Fgf18 mutant alleles identifies unique and redundant roles in skeletal development. Dev. Biol. 411, 72–84.

Jacob, A. L., Smith, C., Partanen, J. and Ornitz, D. M. (2006). Fibroblast growth factor receptor 1 signaling in the osteo-chondrogenic cell lineage regulates sequential steps of osteoblast maturation. Dev. Biol. 296, 315–328.

Kahn, J., Shwartz, Y., Blitz, E., Krief, S., Sharir, A., Breitel, D. A., Rattenbach, R., Relaix, F., Maire, P., Rountree, R. B., et al. (2009). Muscle contraction is necessary to maintain joint progenitor cell fate. Dev. Cell 16, 734–743.

Karuppaiah, K., Yu, K., Lim, J., Chen, J., Smith, C., Long, F. and Ornitz, D. M. (2016). FGF signaling in the osteoprogenitor lineage non-autonomously regulates postnatal chondrocyte proliferation and skeletal growth. Development 143, 1811–1822.

Kim, D., Pertea, G., Trapnell, C., Pimentel, H., Kelley, R. and Salzberg, S. L. (2013). TopHat2: accurate alignment of transcriptomes in the presence of insertions, deletions and gene fusions. Genome Biol 14, R36.

Lanske, B., Karaplis, A. C., Lee, K., Luz, A., Vortkamp, A., Pirro, A., Karperien, M., Defize, L. H. K., Ho, C., Mulligan, R. C., et al. (1996). PTH/PTHrP Receptor in Early Development and Indian Hedgehog--Regulated Bone Growth. Science 273, 663–666.

Leung, V. Y. L., Gao, B., Leung, K. K. H., Melhado, I. G., Wynn, S. L., Au, T. Y. K., Dung, N. W. F., Lau, J. Y. B., Mak, A. C. Y., Chan, D., et al. (2011). SOX9 Governs Differentiation Stage-Specific Gene Expression in Growth Plate Chondrocytes via Direct Concomitant Transactivation and Repression. PLoS Genet 7, e1002356.

Long, F. (2004). Ihh signaling is directly required for the osteoblast lineage in the endochondral skeleton. Development 131, 1309–1318.

Lu, J., Dai, J., Wang, X., Zhang, M., Zhang, P., Sun, H., Zhang, X., Yu, H., Zhang, W., Zhang, L., et al. (2015). Effect of fibroblast growth factor 9 on the osteogenic differentiation of bone marrow stromal stem cells and dental pulp stem cells. Molecular Medicine Reports 11, 1661–1668.

Mau, E., Whetstone, H., Yu, C., Hopyan, S., Wunder, J. S. and Alman, B. A. (2007). PTHrP regulates growth plate chondrocyte differentiation and proliferation in a Gli3 dependent manner utilizing hedgehog ligand dependent and independent mechanisms. Developmental Biology 305, 28–39.

Mead, R. J., Bennett, E. J., Kennerley, A. J., Sharp, P., Sunyach, C., Kasher, P., Berwick, J., Pettmann, B., Battaglia, G., Azzouz, M., et al. (2011). Optimised and Rapid Pre-clinical Screening in the SOD1G93A Transgenic Mouse Model of Amyotrophic Lateral Sclerosis (ALS). PLoS ONE 6, e23244.

Ohbayashi, N., Shibayama, M., Kurotaki, Y., Imanishi, M., Fujimori, T., Itoh, N. and Takada, S. (2002). FGF18 is required for normal cell proliferation and differentiation during osteogenesis and chondrogenesis. Genes Dev. 16, 870–879.

Ornitz, D. M. and Itoh, N. (2015). The Fibroblast Growth Factor signaling pathway. Wiley Interdiscip Rev Dev Biol 4, 215–266.

Ornitz, D. M. and Marie, P. J. (2002). FGF signaling pathways in endochondral and intramembranous bone development and human genetic disease. Genes Dev. 16, 1446–1465.

Ornitz, D. M. and Marie, P. J. (2015). Fibroblast growth factor signaling in skeletal development and disease. Genes Dev. 29, 1463–1486.

Pai, A. C. (1965). Developmental genetics of a lethal mutation, muscular dysgenesis (mdg), in the mouse. Developmental Biology 11, 82–92.

Pan, Y., Bai, C. B., Joyner, A. L. and Wang, B. (2006). Sonic hedgehog Signaling Regulates Gli2 Transcriptional Activity by Suppressing Its Processing and Degradation. Molecular and Cellular Biology 26, 3365–3377.

Peters, K. G., Werner, S., Chen, G. and Williams, L. T. (1992). Two FGF receptor genes are differentially expressed in epithelial and mesenchymal tissues during limb formation and organogenesis in the mouse. Development 114, 233–243.

Rigueur, D. and Lyons, K. M. (2014). Whole-mount skeletal staining. Methods Mol. Biol. 1130, 113–121.

Roberts, R. R., Bobzin, L., Teng, C. S., Pal, D., Tuzon, C. T., Schweitzer, R. and Merrill, A. E. (2019). FGF signaling patterns cell fate at the interface between tendon and bone. Development 146, dev170241.

Robinson, M. D., McCarthy, D. J. and Smyth, G. K. (2010). edgeR: a Bioconductor package for differential expression analysis of digital gene expression data. Bioinformatics 26, 139–140.

Rot-Nikcevic, I., Reddy, T., Downing, K. J., Belliveau, A. C., Hallgrímsson, B., Hall, B. K. and Kablar, B. (2006). Myf5−/−⍰:MyoD−/−amyogenic fetuses reveal the importance of early contraction and static loading by striated muscle in mouse skeletogenesis. Dev. Genes Evol. 216, 1–9.

Sahni, M., Ambrosetti, D. C., Mansukhani, A., Gertner, R., Levy, D. and Basilico, C. (1999). FGF signaling inhibits chondrocyte proliferation and regulates bone development through the STAT-1 pathway. Genes Dev. 13, 1361–1366.

Schneider, C. A., Rasband, W. S. and Eliceiri, K. W. (2012). NIH Image to ImageJ: 25 years of image analysis. Nat. Methods 9, 671–675.

Schwartz, A. G., Long, F., Thomopoulos, S., Karp, S., Gaffield, W., McMahon, A. P. and Vortkamp, A. (2015). Enthesis fibrocartilage cells originate from a population of Hedgehog-responsive cells modulated by the loading environment. Development (Cambridge, England) 142, 196–206.

St-Jacques, B., Hammerschmidt, M. and McMahon, A. P. (1999). Indian hedgehog signaling regulates proliferation and differentiation of chondrocytes and is essential for bone formation. Genes Dev. 13, 2072–2086.

Su, N., Jin, M. and Chen, L. (2014). Role of FGF/FGFR signaling in skeletal development and homeostasis: learning from mouse models. Bone Res 2, 14003.

Tan, Z., Niu, B., Tsang, K. Y., Melhado, I. G., Ohba, S., He, X., Huang, Y., Wang, C., McMahon, A. P., Jauch, R., et al. (2018). Synergistic co-regulation and competition by a SOX9-GLI-FOXA phasic transcriptional network coordinate chondrocyte differentiation transitions. PLoS Genet 14, e1007346.

Tremblay, P., Dietrich, S., Mericskay, M., Schubert, F. R., Li, Z. and Paulin, D. (1998). A crucial role for Pax3 in the development of the hypaxial musculature and the long-range migration of muscle precursors. Dev. Biol. 203, 49–61.

Vaahtokari, A., Aberg, T. and Thesleff, I. (1996). Apoptosis in the developing tooth: association with an embryonic signaling center and suppression by EGF and FGF-4. Development 122, 121–129.

Wang, L., Wang, S. and Li, W. (2012). RSeQC: quality control of RNA-seq experiments. Bioinformatics 28, 2184–2185.

Winkler, D. G., Sutherland, M. K., Geoghegan, J. C., Yu, C., Hayes, T., Skonier, J. E., Shpektor, D., Jonas, M., Kovacevich, B. R., Staehling-Hampton, K., et al. (2003). Osteocyte control of bone formation via sclerostin, a novel BMP antagonist. EMBO J 22, 6267–6276.

Yang, Y. and Kozin, S. H. (2009). Cell signaling regulation of vertebrate limb growth and patterning. J Bone Joint Surg Am 91 Suppl 4, 76–80.

Zelzer, E., Blitz, E., Killian, M. L. and Thomopoulos, S. (2014). Tendon-to-bone attachment: from development to maturity. Birth Defects Res. C Embryo Today 102, 101–112.

